# The composition of northeast pacific fishes in a fish tank examined by eDNA metabarcoding

**DOI:** 10.1101/2020.12.21.423745

**Authors:** Sergei V. Turanov, Olesia A. Rutenko

## Abstract

The taxonomy of fish in the northeast Pacific area has been recently revised using molecular genetic methods, including the development of a reference library of DNA fragments for species identification. Such libraries are the basis for the development of non-invasive, high-throughput methods for monitoring biodiversity using environmental DNA (eDNA). In order to validate this approach, we used a water eDNA metabarcoding technique based on *12S rRNA* and *COI* mitochondrial fragments and assessed the composition of the twenty northeast Pacific fish species held in a fish tank at the Primorsky Aquarium (Vladivostok, Russia). Only the *12S* fragment revealed data on fish-related operational taxonomic units (OTUs). Approximately 68% of the reads were classified into two species of the genus *Oncorhynchus*, whose shredded fillet is used for feeding. According to the taxonomic identification for the rest of the reads, 8 out of 20 fish species in the tank (40%) were identified unambiguously, while two species could not be identified. Ten taxa can be considered conditionally identifiable since they might be concealed behind a conflicting taxonomic identification at the genus or family level. In this case, an improvement of the reference library would provide resolution. We detected contamination, which may be related to both intra-laboratory contaminants occurring during DNA extraction and water intake supplying the fish tank.

## Introduction

Species diversity of fish and fish-like vertebrates of Russian waters includes more than 1400 species (Parin et al. 2014, Orlov and Tokranov 2019), of which more than 1300 inhabit sea waters. Seas of the northeast Pacific Ocean (Sea of Japan, Sea of Okhotsk and the Bering Sea) are characterized by a significant diversity of fish taxa, the systematics of which have been recently revised with increasing use of molecular genetic approaches (Radchenko 2015, Moreva et al. 2017, 2019, Orlova et al. 2019, Turanov et al. 2019, Balakirev et al. 2020, Radchenko et al. 2020). Unified methods are also used in this area, which aim to create a reference library of DNA fragments for accurate fish identification (Mecklenburg, Møller, & Steinke, 2011; Turanov et al., 2016; Zhang & Hanner, 2011). This strategy also provides a foundation for the development of high-throughput non-invasive diversity monitoring methods based on environmental DNA (McGee et al. 2019, Weigand et al. 2019, Schenekar et al. 2020). Environmental DNA (eDNA) uses extracellular DNA molecules that are spread by any organism regenerating its own tissues over time, and can be used to trace that organism through environmental samples (Taberlet et al. 2018). Indeed, DNA from the aquatic environment is widely used to assess taxonomic diversity (Thomsen et al. 2012). It can also be used to approximate fish biomass estimates (Salter et al, 2019; Takahara et al., 2012) and provide information on genetic diversity in terms of haplotype variation (Tsuji et al. 2020). However, this approach is not particularly popular in Russia, and it applies exclusively to the aquatic biodiversity in the west of the country (Velichko et al. 2014, Kirilchik 2018, Lecaudey et al. 2019, Milyutina et al. 2019, Belevich et al. 2020). In order to validate these methods for non-invasive monitoring, mock community analysis (Braukmann et al. 2019) has been used; with respect to fish, taxonomic diversity in aquarium with known fish composition has been analyzed (Kelly et al. 2014, Morey et al. 2020). This makes it possible to assess potential limitations of the approach and generate the necessary methodological recommendations when analyzing natural samples. The present paper reports the evaluation of the northeastern Pacific marine fish composition in a fish tank in the Primorsky Aquarium using eDNA metabarcoding techniques.

## Materials and methods

Water from a tank (volume 1.68 m^3^) at the Primorsky Aquarium (Russky Island, Vladivostok, Primorsky Krai, Russia) was taken on December 12, 2019. This tank holds representatives of the coast of the Far Eastern sea area (Sea of Japan, Sea of Okhotsk, and the Bering Sea). The water temperature is maintained in the range of 10–13□C. Taxonomic information and the number of fishes is provided in the Appendix (Suppl. material 1: Table S1). Fishes in this tank are fed with minced fillet of pink (*Oncorhynchus gorbuscha* (Walbaum, 1792)) and chum salmons (*O. keta* (Walbaum, 1792)), greenlings (*Hexagrammos* sp. Tilesius, 1810), chub mackerel (*Scomber japonicus* Linnaeus, 1758), squids (*Todarodes pacificus* (Steenstrup, 1880), *Berryteuthis magister* (Berry, 1913)), and shrimp (*Pandalus latirostris Rathbun, 1902*). Three replicate samples of water (about 450 ml each) were collected from the tank with a syringe (150 ml capacity, luer lock type). The entire volume of water in each replicate was pushed through a single syringe filter (diameter: 25 mm, pore size: 0.45 μm, material: PES). The DNA on the filter was fixed by pushing 1 ml of Longmire’s buffer through the filter, and next the inlet and outlet holes were closed with combi-stopper plugs. The fixed filter was then stored at −2MC. DNA was extracted from the pre-fixed filter using the M-Sorb-OOM kit (Sintol, Moscow). The manufacturer’s protocol was modified such that the lysis buffer was heated to 65 □C, pushed in the reverse direction of filtration (Kesberg and Schleheck 2013) and poured into a clean test tube. The isolated DNA was stored at −2MC. Two mitochondrial fragments - *12S rRNA* with a length of 170 bp (Miya et al. 2015) and *COI* with a length of 320 bp (Geller et al. 2013, Leray et al. 2013, Wangensteen et al. 2018) - were used to retrieve the information on the taxonomic diversity from the eDNA samples. A pair of primers with an individual 6-nucleotide tag (doubly-tagged) developed in ecotag (Boyer et al. 2016) was used to amplify the fragments for each sample. Negative PCR controls for each fragment were also performed by using separate pairs of tagged primers. The PCR for each sample was performed using three replicates. The reaction mixture included 10 μl of AmpliTaq Gold 360 Master Mix (Thermofisher), 0.5 μl of each of the forward and reverse primers (10 μM), 0.16 μl of bovine serum albumin, 10 ng of DNA and deionized water to the final volume (20 μl). The thermal cycling profile for the amplification of the *COI* fragment included preheating at 95 □C for 10 minutes, followed by 35 cycles according to the following scheme: 1 min. at 94 □C, 1 min. at 45 □C and 1 min. at 7MC. The final elongation was at 7MC for 5 min. The *12S rRNA* fragment was amplified as follows: preheating at 95□C for 3 min., 35 cycles of 9MC for 20 sec, 60□C for 15 sec, and 7MC for 15 sec., and a final elongation step at 7MC for 5 min. The amplification results were examined by running the fragments on a 1% agarose gel followed by exposure in ethidium bromide solution and visualization under ultraviolet light. The amplicons were purified with Cleanup S-Cap (Evrogen, Moscow) and normalized (see (Elbrecht and Steinke 2019)) before pooling. The amount of control reactions was taken as the average from the resulting volume of the normalized samples before pooling. Next, the normalized amplicons were combined with the PCR controls and sequenced at Novogene. The library was generated using NEBNext Ultra II DNA Library Prep Kit for Illumina (New England Biolabs, England) and sequenced with an Illumina high-throughput sequencer using a 250 bp paired-end sequencing strategy.

The obtained reads were processed according to the Begum metabarcoding pipeline (Zepeda-Mendoza et al. 2016, Yang et al. 2020). After adapter removal and preliminary evaluation of the read quality with fastqc, possible read errors were corrected in Spades (Bankevich et al. 2012). Next, paired reads were merged into consensus sequences using PandaSeq (Masella et al. 2012). We then used Begum to demultiplex and filter the reads for each sample. Clustering was done by sumaclust (Mercier et al. 2013) with parameters -t 0.98 and -R 0.85. The taxonomic assignment for the generated operational taxonomic units (OTUs) was performed using BLAST (Camacho et al. 2009) implemented in Ubuntu command line interface. The taxonomic information was then summarized in a single table based on the output from the MEGAN community edition program (Huson et al. 2016). The top percent of LCA parameters was set to 2.0 implementing the naïve LCA algorithm with 80 percent to cover. The number of OTUs after taxonomic referencing was corrected using the lulu package (Frøslev et al. 2017) with the following parameters: minimum match - 95%, minimum relative cooccurrence - 0.97.

## Results and discussion

After demultiplexing, the *12S rRNA* fragment had 42260 reads, 2827 of which were unique. The *COI* fragment accounted for 591 sequences, of which 241 were unique. The controls identified 60 reads for the *12S* fragment and no reads for *COI*. After filtering the reads and deleting the chimeric sequences, there were 38441 sequences for *12S rRNA* and 289 sequences for *COI*. The clustering of *COI* sequences revealed 13 OTUs (Suppl. material 2: Table S2). Most of the reads (289) with six OTUs were not identified (not classified to any of the life domains). Two OTUs with 12 reads in total were assigned to the species *Pedobacter ginsengisoli* and *Polaribacter* sp. The remaining five OTUs were classified as eukaryotes, four of which belonged to different multicellular organisms from Annelida (*Polydora* sp.), Arthropoda (family Miraciidae) and Cnidaria (genus *Clytia*); one OTU belonged to *Phytophthora pseudosyringae* from the clade SAR.

Sequences of *12S rRNA* clustered into 59 OTUs, four of which were assigned to bacteria with 57 reads in total. The remaining 55 OTUs were related to ray-finned fishes, classified into seven orders, 15 families and 28 genera (Fig. 1). Species-level taxonomy was unambiguously assigned to 23 OTUs (39%). There were 15 OTUs (25%) with taxonomic contradictions at the species level; at the genus level, there were nine OTUs (15%), and one OTU at the family level. The conflict in this case is defined as equal (or undetectable) probability of assigning the same OTU to different taxa according to BLAST. This may indicate either erroneous taxonomic identification of one or several sequences that make up the reference library, or that the marker is highly conserved hence divergence is not sufficient enough to distinguish two or more species. These and other limitations of the approach have been mentioned earlier when estimating fish diversity (Collins et al. 2019, Weigand et al. 2019, Schenekar et al. 2020, Stoeckle et al. 2020). On the other hand, despite the data correction with the lulu R package, some species names were simultaneously assigned to two (*Hypsagonus jordani, Ulcina olrikii*) and even three (*Hypomesus nipponensis, Oncorhynchus keta, Opisthocentrus ocellatus*) OTUs (see Suppl. material 3: Table S3). Selecting a standard delineation threshold for clustering based on hierarchical approaches can distort the true number of OTUs. Arguably, more accurate results can be expected from clustering approaches that are based on the natural organization of data without setting a hard cut-off threshold (Hao et al. 2011).

**Figure 1.**
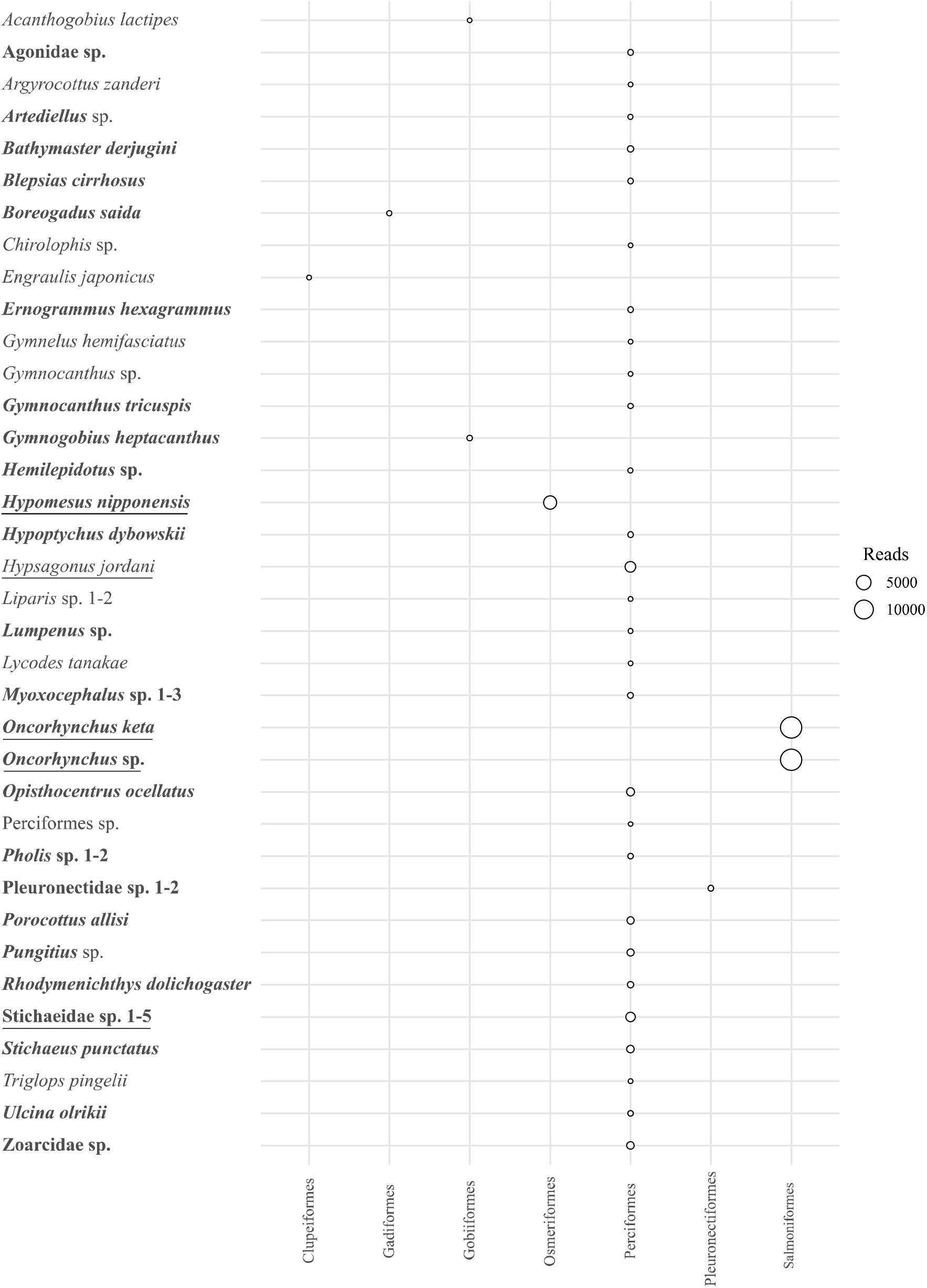
A bubble graph showing the number of reads for different OTUs obtained for *12S rRNA*, amplified based on the water eDNA samples from a fish tank at the Primorsky Aquarium. The orders are presented horizontally and the OTUs are placed vertically. The size of circles demonstrates the number of reads assigned to the different OTUs. OTU names in **bold** have no less than 20 reads. Each <u>underlined</u> OTU accounts for more than 1,000 reads.

The largest number of reads came from two species of the genus *Oncorhynchus* (whose fillets are used for feeding, see above) – more than 13000 for each species. Hence, it is encouraged to account for this when planning sufficient sequencing depth for mesocosm experiments (Kelly et al. 2014, Morey et al. 2020) to provide adequate coverage for the detection of DNA for targeted organisms with low amounts of eDNA. An option would be to use blocking primers for the forage organisms, similar to those used in diet assessment. However, it is necessary to consider all the advantages and limitations of this approach (Piñol et al., 2015). Primer bias should not be excluded either (Clarke et al. 2014, Deiner et al. 2017), although its impact in this case seems to be less likely. More than 1000 reads were accounted for each of the clusters among *Hypomesus nipponensis* McAllister, 1963, *Hypsagonus jordani* Jordan & Starks, 1904 and Stichaeidae sp. - 26 OTUs had 20 to 1200 reads, whereas each of the other 23 OTUs had less than 20 reads.

We unambiguously identified eight out of 20 taxa (40%) in the tank. Two taxa present (*Alcichthys elongatus* (Steindachner 1881), *Bero elegans* (Steindachner, 1881)) have not been identified, i.e. there is no OTU that might be related to these species. Another 10 taxa (50%) may somehow be concealed behind OTUs with conflicting taxonomic reference (see above), such as *Chirolophis* sp., *Lumpenus* sp., *Pungitius* sp., Pleuronectidae sp. (Fig. 1, Suppl. material 3: Table S3). The indication of all these taxa except for *Aspidophoroides* Lacepède, 1801 and *Boreogadus* Günther, 1862 in the aquarium may come from environmental DNA originating from the water storage reservoir, whose intake is located next to the Primorsky Aquarium building. Hence, these OTUs may be the result of contamination during DNA isolation or at the PCR stage. The potential for this type of contamination, as well as the presence of hidden sources of environmental DNA, is expected and, regretfully, is not unusual (Kelly et al. 2014, Morey et al. 2020). In addition to reducing the probability of contamination, the development and verification of a reference library for fish identification should be considered as an important way to reduce uncertainty and improve the reliability of high-throughput monitoring methods based on aquatic eDNA (Collins et al. 2019, Weigand et al. 2019).

This preliminary study demonstrates the possible methodological bias of high-throughput DNA-based monitoring from the aquatic environment, and will serve as a starting point for the efficient planning of surveys employing non-invasive methods to monitor aquatic biological diversity of fish in the Russian Far East.

## Acknowledgements

This research was partially supported by a Grant of the President of the Russian Federation (MK-305.2019.4) and Ministry of Science and Higher Education of the Russian Federation (agreement number 075-15-2020-796, grant number 13.1902.21.0012).

## Appendices

**Supplementary material 1, Table S1**. Fish species composition in the tank of the Primorsky Aquarium where marine aquatic eDNA was collected from. Unambiguously identified taxa are highlighted by blue color. Species that were not identified are highlighted by red color.

**Supplementary material 2, Table S2**. Assigned OTUs for *COI* marker.

**Supplementary material 3, Table S3**. Assigned OTUs for *12 rRNA* marker.

