## Supplementary material for "The composition of northeast pacific fishes in a fish tank examined by eDNA metabarcoding": Table S1

**Table S1**. Fish species composition in the tank of the Primorsky Aquarium where marine aquatic eDNA was collected from. Unambiguously identified taxa are highlighted by blue color. Species that were not identified are highlighted by red color.

| **No** | **Species** | **Number of specimens** |
| --- | --- | --- |
| 1 | *Agonomalus proboscidalis* | 1 |
| 2 | *Alcichthys elongatus* | 3 |
| 3 | *Bathymaster derjugini* | 1 |
| 4 | *Bero elegans* | 3 |
| 5 | *Blepsias cirhosus* | 1 |
| 6 | *Brachyopsis segaliensis* | 2 |
| 7 | *Chirolophis japonicus* | 3 |
| 8 | *Chirolophis saitone* | 2 |
| 9 | *Gymnocanthus herzensteini* | 5 |
| 10 | *Gymnogobius heptacanthus* | 5 |
| 11 | *Hypomesus japonicus* | 3 |
| 12 | *Hypsagonus jordani* | 2 |
| 13 | *Hypoptychus dybowskii* | 5 |
| 14 | *Lumpenus sagitta* | 2 |
| 15 | *Opisthocentrus ocellatus* | 3 |
| 16 | *Pholidapus dybowskii* | 10 |
| 17 | *Podothecus gilberti* | 3 |
| 18 | *Pseudopleuronectes* sp. | 2 |
| 19 | *Pungitius* sp. | 6 |
| 20 | *Stichaeopsis epallax* | 5 |
